# Human Papillomavirus 16 replication converts SAMHD1 into a homologous recombination factor and promotes its recruitment to replicating viral DNA

**DOI:** 10.1101/2023.11.13.566899

**Authors:** Claire D. James, Aya Youssef, Apurva T. Prabhakar, Raymonde Otoa, Austin Witt, Rachel L. Lewis, Molly L. Bristol, Xu Wang, Kun Zhang, Renfeng Li, Iain M. Morgan

**Affiliations:** Virginia Commonwealth University (VCU), Philips Institute for Oral Health Research, School of Dentistry, Richmond, VA, 23298; VCU Massey Cancer Center, Richmond, VA, 23298; Department of Microbiology and Molecular Genetics, University of Pittsburgh, PA, 15232; Hillman Cancer Center, University of Pittsburgh Medical Center, PA, 15232

**Keywords:** human papillomavirus, replication, life cycle, SAMHD1, cervical cancer, head and neck cancer, therapy, homologous recombination, phosphorylation

## Abstract

We have demonstrated that SAMHD1 (sterile alpha motif and histidine-aspartic domain HD-containing protein 1) is a restriction factor for the HPV16 life cycle. Here we demonstrate that in HPV negative cervical cancer C33a cells and human foreskin keratinocytes immortalized by HPV16 (HFK+HPV16), SAMHD1 is recruited to E1-E2 replicating DNA. Homologous recombination (HR) factors are required for HPV16 replication and viral replication promotes phosphorylation of SAMHD1, which converts it from a dNTPase to an HR factor independent from E6/E7 expression. A SAMHD1 phosphor-mimic (SAMHD1 T592D) reduces E1-E2 mediated DNA replication in C33a cells and has enhanced recruitment to the replicating DNA. In HFK+HPV16 cells SAMHD1 T592D is recruited to the viral DNA and attenuates cellular growth, but does not attenuate growth in isogenic HFK cells immortalized by E6/E7 alone. SAMHD1 T592D also attenuates the development of viral replication foci following keratinocyte differentiation. The results indicated that enhanced SAMHD1 phosphorylation could be therapeutically beneficial in cells with HPV16 replicating genomes. Protein phosphatase 2A (PP2A) can dephosphorylate SAMHD1 and PP2A function can be inhibited by endothall. We demonstrate that endothall reduces E1-E2 replication and promotes SAMHD1 recruitment to E1-E2 replicating DNA, mimicking the SAMHD1 T592D phenotypes. Finally, we demonstrate that in head and neck cancer cell lines with HPV16 episomal genomes endothall attenuates their growth and promotes recruitment of SAMHD1 to the viral genome. The results suggest that targeting cellular phosphatases has therapeutic potential for the treatment of HPV infections and cancers.

**Importance:** Human papillomaviruses are causative agents in around 5% of all human cancers. The development of anti-viral therapeutics depends upon an increased understanding of the viral life cycle. Here we demonstrate that HPV16 replication converts SAMHD1 into an HR factor via phosphorylation. This phosphorylation promotes recruitment of SAMHD1 to viral DNA to assist with replication. A SAMHD1 mutant that mimics phosphorylation is hyper-recruited to viral DNA and attenuates viral replication. Expression of this mutant in HPV16 immortalized cells attenuates the growth of these cells, but not cells immortalized by the viral oncogenes E6/E7 alone. Finally, we demonstrate that the phosphatase inhibitor endothall promotes hyper-recruitment of endogenous SAMHD1 to HPV16 replicating DNA and can attenuate the growth of both HPV16 immortalized human foreskin keratinocytes and HPV16 positive head and neck cancer cell lines. We propose that phosphatase inhibitors represent a novel tool for combating HPV infections and disease.

## Introduction

Human papillomaviruses infect epithelial cells and cause around 5% of all cancers (1). During the viral life cycle, the virus exists as an 8 kbp double stranded DNA episome replicated by two viral factors, E1 and E2, in association with host factors (2-6). The process of viral replication activates the DNA damage response (DDR) and recruits a host of DNA repair factors to the viral DNA, including TopBP1, SIRT1 and WRN (7-15). Activation of the DDR is critical for the HPV life cycle, as is recruitment and interaction with host DNA repair factors, particularly those involved in homologous recombination (12, 13, 16-21). It is proposed that the recruitment of host DDR factors to the viral genome promote homologous recombination (HR) mediated DNA replication during the viral life cycle (22). Our understanding of the host factors involved in promoting HR mediated replication of HPV genomes remains incomplete, and identification of such factors will potentially facilitate the identification of novel therapeutic approaches for disrupting viral replication and treating HPV related diseases.

SAMHD1 regulates intracellular levels of dNTPs using triphosphohydrolase (dNTPase) enzyme activity (23). dNTPase function depends upon formation of a homo-tetramer and each SAMHD1 molecule consists of a sterile alpha motif (SAM) that can interact with other SAM domains, and a dGTP-regulated dNTP hydrolase domain (HD) that regulates the dNTP levels in cells (24-26). SAMHD1 can act as a restriction factor for several viruses, including HIV (27-29). SAMHD1 can also modulate cytomegalovirus infection by controlling NF-kB activity (30). Viruses can also counteract SAMHD1 function: HIV-2 Vpx promotes proteasomal degradation of SAMHD1 (31); conserved herpesvirus protein kinases can antagonize SAMHD1 restriction via phosphorylation (32). For HPV16, deletion of SAMHD1 expression using CRISPR targeting generated several phenotypes, including increased cellular proliferation, increased DNA damage, and increase viral genome amplification (33). These phenotypes indicate that SAMHD1 also acts as a restriction factor for HPV16. However, the mechanism of how SAMHD1 acts as an HPV16 restriction factor remained unresolved.

The phosphorylation of SAMHD1 on threonine 592 (T592) by CDK1 or CDK2 converts SAMHD1 into a homologous recombination factor promoting the recruitment of MRE11 to sites of damaged DNA for end resection (34, 35). The binding to DNA for end resection is promoted by deacetylation of SAMHD1 by SIRT1, a class III deacetylase (36). SIRT1 can deacetylate HPV16 E2 to promote stability and boost E2 replication function (8). The phosphorylation of SAMHD1 has been proposed to reduce dNTPase activity as it destabilizes SAMHD1 homo-tetramerization (23), although others have suggested that the dNTPase activity is not affected by phosphorylation (28). Mechanistically, if phosphorylation did disrupt the dNTPase activity and promote the HR function of SAMHD1 to repair DNA, it would both boost HR as well as increase the nucleotide pool available for DNA replication/repair.

Given the role of HR in the HPV16 life cycle and the recruitment of HR factors to the viral genome to promote viral replication, this report investigated the mechanistic role of SAMHD1 in the HPV16 life cycle. The results demonstrate that SAMHD1 is recruited to HPV16 E1-E2 replicating DNA, both in C33a viral replication models and in human foreskin keratinocytes immortalized by HPV16 (HFK+HPV16). SAMHD1 is phosphorylated on T592 only by the full HPV16 genome, the viral oncogenes cannot do this, suggesting that viral replication *per se* induces the phosphorylation. A SAMHD1 phosphor-mimetic (T592D, where the threonine is mutated to an aspartic acid) is hyper-recruited to E1-E2 replicating DNA and results in an attenuation of viral replication levels. In HFK+HPV16 cells, SAMHD1 T592D attenuates the growth of the cells and prevents the formation of viral replication foci that are induced in differentiating HFK+HPV16 cells. The results suggest a model in which increasing SAMHD1 recruitment to viral DNA by increasing phosphorylation disrupts viral replication, along with host replication, and attenuates the growth of HPV16 positive cells. SAMHD1 is dephosphorylated by PP2A during mitotic exit (37), and endothall is an inhibitor of PP2A (38). Endothall treatment attenuated E1/E2 replication in C33a cells and enhanced recruitment of SAMHD1, E1 and E2 to the replicating DNA, mimicking the effects of SAMHD1 T592D. HFK+HPV16 treatment with endothall enhanced the recruitment of SAMHD1 and E2 to the viral genomes and attenuated cellular growth as well as inhibiting the development of replication foci following differentiation. All of these results mimicked the results generated with SAMHD1 T592D. Strikingly, the effects of endothall and SAMHD1 T592D on cellular growth were dependent upon the presence of the full-length replicating HPV16 genome, endothall had minimal effects on the growth of isogenic cells immortalized by the HPV16 oncogenes E6 and E7. Finally, we demonstrate that the head and neck cancer cell line, SCC-104, which contains episomal HPV16 genomes, is hypersensitive to endothall and treatment resulted in increased SAMHD1 and E2 recruitment to the replicating HPV16 genomes. Endothall was less toxic to head and neck cancer cell lines with integrated HPV16 genomes (SCC-47) or an HPV negative head and neck cancer cell line (HN30). We propose that phosphatase inhibitors represent a novel approach for the treatment of HPV16 positive tumors that have replicating HPV16 genomes.

## Results

### HPV16 replication phosphorylates SAMHD1 to convert it to a homologous recombination factor, and recruits it to E1-E2 replicating DNA

We established that E1-E2 replicating DNA recruits SAMHD1 using our C33a model system (10, 39). SAMHD1 levels were knocked down using an shRNA in the presence of E1 and E2 expression along with a plasmid containing the origin of replication (Figure 1A, compare lane 4 with lane 3). Knockdown of SAMHD1 did not alter the levels of E1 or E2 protein expression. Chromatin immunoprecipitation experiments using chromatin prepared from C33a cells transfected with E1-E2-pOri and shRNA Control (Ctrl) or shRNA SAMHD1 demonstrated that the knockdown of SAMHD1 expression (Figure 1A) resulted in a reduction of SAMHD1 signal (Figure 1B, compare lane 6 with lane 3). In the absence of E1 or E2 there was minimal ChIP signal detected as reported previously (not shown, (7, 8, 10, 39-42)). The knockdown of SAMHD1 expression did not significantly alter E1-E2 DNA replication levels in C33a cells (Figure 1C). Next, we determined that SAMHD1 was recruited to viral genomes in human foreskin keratinocytes immortalized by HPV16 (HFK+HPV16) (Figure 1D). Chromatin was prepared from two independent donor HFK+HPV16 cell lines (21) and ChIP assays carried out in triplicate with HA, E2 and SAMHD1 antibodies. The levels of HPV16 LCR DNA pulled down were determined relative to input, expressed as percentages, and represent a summary of three independent experiments. The HA antibody pulls down background levels of LCR (lanes 1&2) while both E2 (lanes 3&4) and SAMHD1 (lanes 5&6) pulled down significantly more LCR DNA in both donor lines. The results suggested that SAMHD1 was converted into an HR factor in HFK+HPV16 and this was confirmed by determining the phosphorylation status of SAMHD1 (Figure 1E). HFKs from two different donors were immortalized by hTERT, the full HPV16 genome (WT HPV16) or E6 and E7 expression via retroviral transduction (E6E7). SAMHD1 levels were similar in all immortalized lines. However, an antibody specific for SAMHD1 phosphorylated on threonine 592 (SAMHD1 p592, middle panel) detected SAMHD1 phosphorylation only in the presence of the entire HPV16 genome (lanes 2 and 5). It is likely that the DDR induced by HPV16 replication drives the phosphorylation of SAMHD1, and this is not due to E6/E7 expression (lanes 3 and 6). Figure 1 demonstrates that E1 and E2 replicating DNA recruits SAMHD1, and that HPV16 full genome converts SAMHD1 into an HR factor.

**Figure 1.**
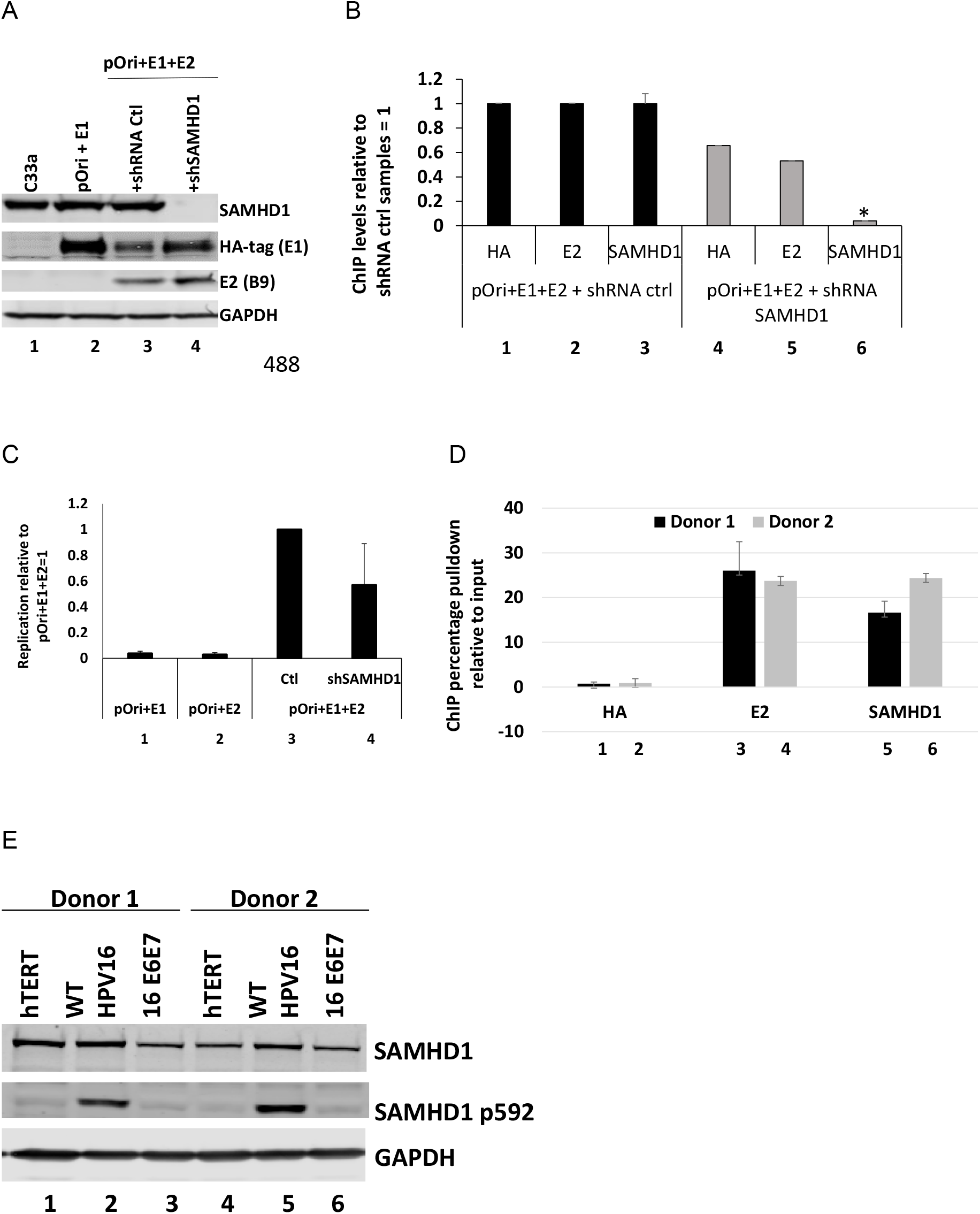
A. C33a cells were transfected with the indicated plasmids, and the expression levels of E2, E1 (HA tagged) and SAMHD1 determined. B. Chromatin was prepared from the indicated transfected cells and ChIP assays carried out to detect the levels of E1 (HA), E2 and SAMHD1 protein recruited to the replicating DNA. There was a significant (*, p<0.05)) reduction in SAMHD1 recruitment when the protein was knocked down, demonstrating that the signal detected is due to SAMHD1 recruitment to the E1-E2 replicating DNA. C. Quantitative replication assays demonstrate that the knockdown of SAMHD1 does not significantly change E1-E2 DNA replication levels. D. Chromatin was prepared from HFK+HPV16 (human foreskin keratinocytes immortalized by HPV16) cells and the presence of E2 and SAMHD1 on the viral genome confirmed using ChIP assays. The HA antibody serves as a negative control and represents background levels; both the E2 and SAMHD1 antibodies significantly increased the signal over background. E. Western blotting of protein extracts from the indicated cell lines demonstrate that SAMHD1 is phosphorylated (pSAMHD1) only in the presence of the full HPV16 genome.

### SAMHD1 is in a cellular complex with E1-E2 and a SAMHD1 phosphor-mimetic is hyper-recruited to E1-E2 replicating DNA, disrupting viral replication

Phosphatases are integral to the control of mammalian DNA replication (43). Figure 1E demonstrated that HPV16 replication converts SAMHD1 into an HR factor via threonine 592 phosphorylation, promoting recruitment to the viral genome. We hypothesized that disrupting de-phosphorylation of SAMHD1 may block viral replication by preventing removal of SAMHD1 from the replication complex. To investigate this, we used SAMHD1 T592 mutants (T592D, threonine to aspartic acid mimicking phosphorylation; T592A, threonine to alanine mimicking non-phosphorylation). Figure 2A demonstrates that the expression of wild-type and SAMHD1 mutants did not significantly alter the expression of the viral proteins E1 and E2 (lanes 7-9, input). The HA-IP (middle panel) immunoprecipitates the HA-tagged E1; E1 interacts with endogenous SAMHD1 (lane 3) and the three SAMHD1 proteins that are overexpressed (lanes 7-9). As expected, E1 interacts with E2 and this is not disrupted by overexpression of any of the SAMHD1 proteins (lanes 5-9). E2 also interacts with endogenous SAMHD1 (bottom panel, lane 4) and the three overexpressed SAMHD1 proteins (lanes 7-9). E2 also interacted with E1 (lanes 5-9), as expected. Replication assays with the overexpressed SAMHD1 proteins demonstrates that SAMHD1 T592D significantly reduced replication (Figure 2B, lane 5). E1 plasmid DNA levels detected in the cell extracts were not significantly altered by the overexpression of the SAMHD1 proteins, demonstrating that they were not toxic to the transfected cells (Figure 2B). ChIP assays (Figure 2C) demonstrated that the expression of SAMHD1 T592D resulted in a significant increase of E1(HA), E2 and SAMHD1 T592D (V5) recruitment to replicating DNA when compared with overexpression of SAMHD1 WT or SAMHD1 T592A (lanes 4, 8 and 12). The overexpression of the latter two proteins did not alter the recruitment of E1 or E2 to replicating DNA, when compared with no SAMHD1 over expression control (pLX304). These results supported our hypothesis that a failure to de-phosphorylate SAMHD1 (T592D mimics “permanent” phosphorylation) can block E1-E2 mediated DNA replication by “freezing” the replication complex on the DNA.

**Figure 2.**
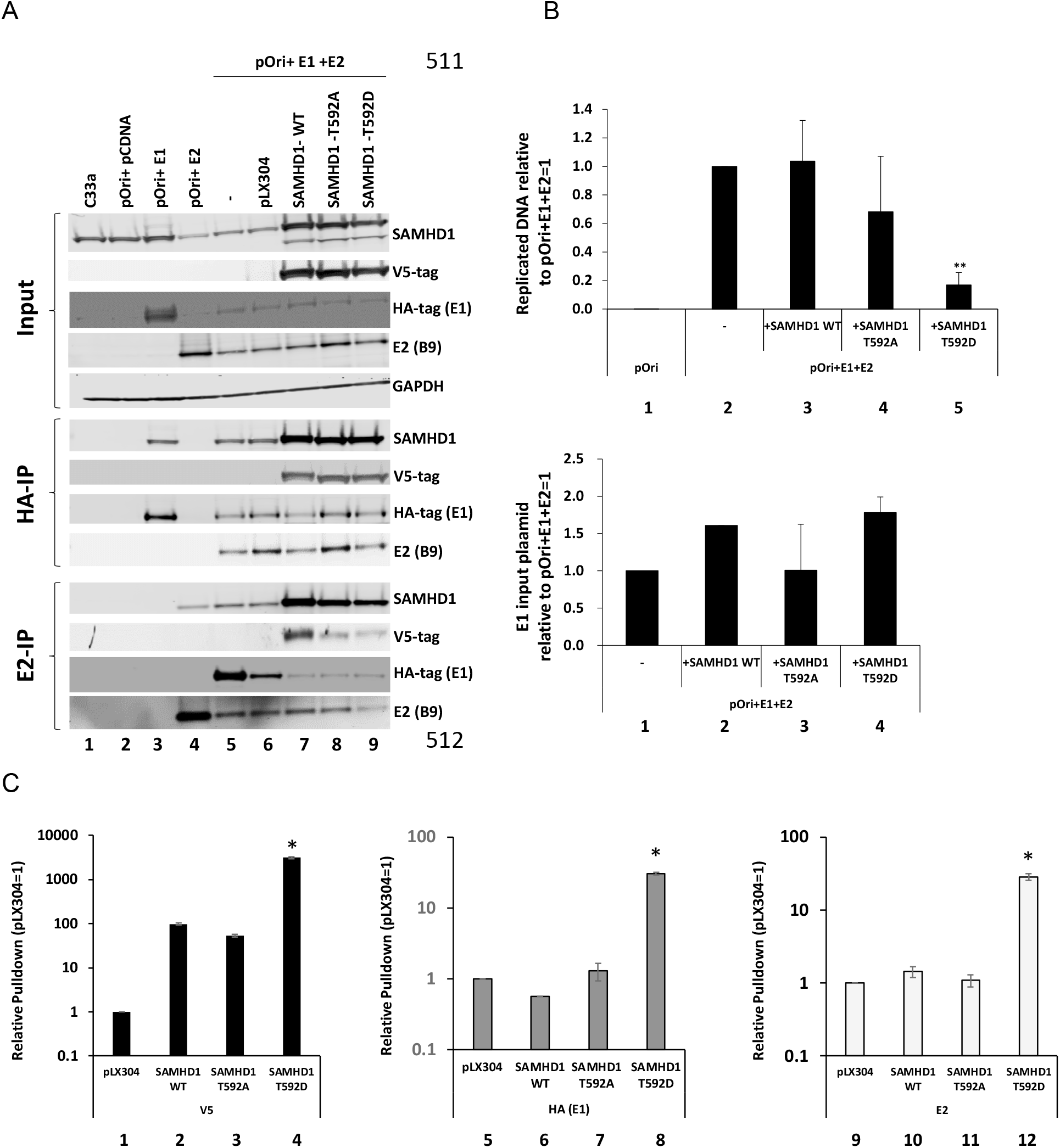
A. C33a cells were transfected with the indicated plasmids, and protein extracts prepared. Input levels of the proteins under study are shown in the top panels (Input). HA (E1) immunoprecipitation demonstrates that E1 can interact with E2 and endogenous SAMHD1 (lanes 3 and 5) as well as with the three co-transfected SAMHD1 expression vector proteins (lanes 7-9). Similarly, E2 can interact with E1 and endogenous SAMHD1 (lanes 4 and 5) and with the three co-transfected SAMHD1 expression vector proteins (lanes 7-9). B. The expression of SAMHD1 T592D significantly (**, p-value < 0.05) reduced E1-E2 DNA replication levels in C33a cells (top panel, compare lanes 2 and 5). The bottom panel monitors the levels of E1 plasmid (that is not replicated during the assay) and there is no significant difference in the detection of this DNA between any of the samples that were transfected with the E1 plasmid. This demonstrates that the reduction of replication with SAMHD1 T592D is not due to cellular toxicity of the protein killing the transfected cells. C. Chromatin was prepared from the transfected cells and ChIP assays carried out with V5 (detects the SAMHD1 proteins from the co-transfected plasmids), HA (detects E1) and E2 antibodies. For all proteins, there is a significant (*, p-value < 0.05) increase in signal in the presence of SAMHD1 T592D. Please note the log scale in this figure.

To investigate the relevance of the results in Figure 2 to the viral life cycle, SAMHD1 WT and mutants were stably over-expressed in HFK+HPV16 cells using lentiviral delivery and selection. Figure 3A demonstrates expression of the SAMHD1 proteins from the delivered lentiviruses in HFK+HPV16 (lanes 6-8 and 14-15) and HFK+E6/E7 (lanes 2-4 and 10-12) in two different foreskin donor lines. ChIP assays targeting the HPV16 LCR were carried out with the HPV16+HFK cells and the percent of input chromatin pulled down determined using E2, V5 (to detect the exogenously overexpressed SAMHD1 proteins) and a control HA antibody (Figure 3B). The results with the HA antibody (lanes 17-24) detected background levels of LCR pulled down. E2 pulled down significantly more LCR DNA than the HA antibody, there was no significant difference in the E2 signal between any of the samples (lanes 1-8). All of the V5 tagged SAMHD1 proteins were recruited to the viral DNA, although there were significant differences between them. SAMHD1 WT was recruited and the SAMHD1 T592A mutant was significantly less recruited (compare lanes 10 and 14 with 11 and 15, respectively). SAMHD1 T592D was the most recruited to the viral DNA and was significantly more than SAMHD1 WT and SAMHD1 T592A (compare lanes 12 and 16 with 10,11 and 14,15). The results support the hypothesis that phosphorylation of SAMHD1 on T592 promotes recruitment of the protein to replicating viral DNA. The viral DNA levels in the cells were not significantly different between all of the samples (not shown), however cellular growth was attenuated by overexpression of the SAMHD1 T592D protein (Figure 3C). In HFK+E6E7 expression of the SAMHD1 proteins did not alter cell growth, in HFK+HPV16 cells growth was attenuated only by the SAMHD1 T592D. The results in Figure 3 suggest that over recruitment of SAMHD1 (T592D) to replicating viral DNA attenuates cellular growth. As viral replication induces a DNA damage response, which would convert SAMHD1 into an HR factor, SAMHD1 T592D could attenuate host cell replication and growth due to the active DNA damage response requiring SAMHD1 recruitment to, and disengagement from, host replication sites to manage cell growth.

**Figure 3.**
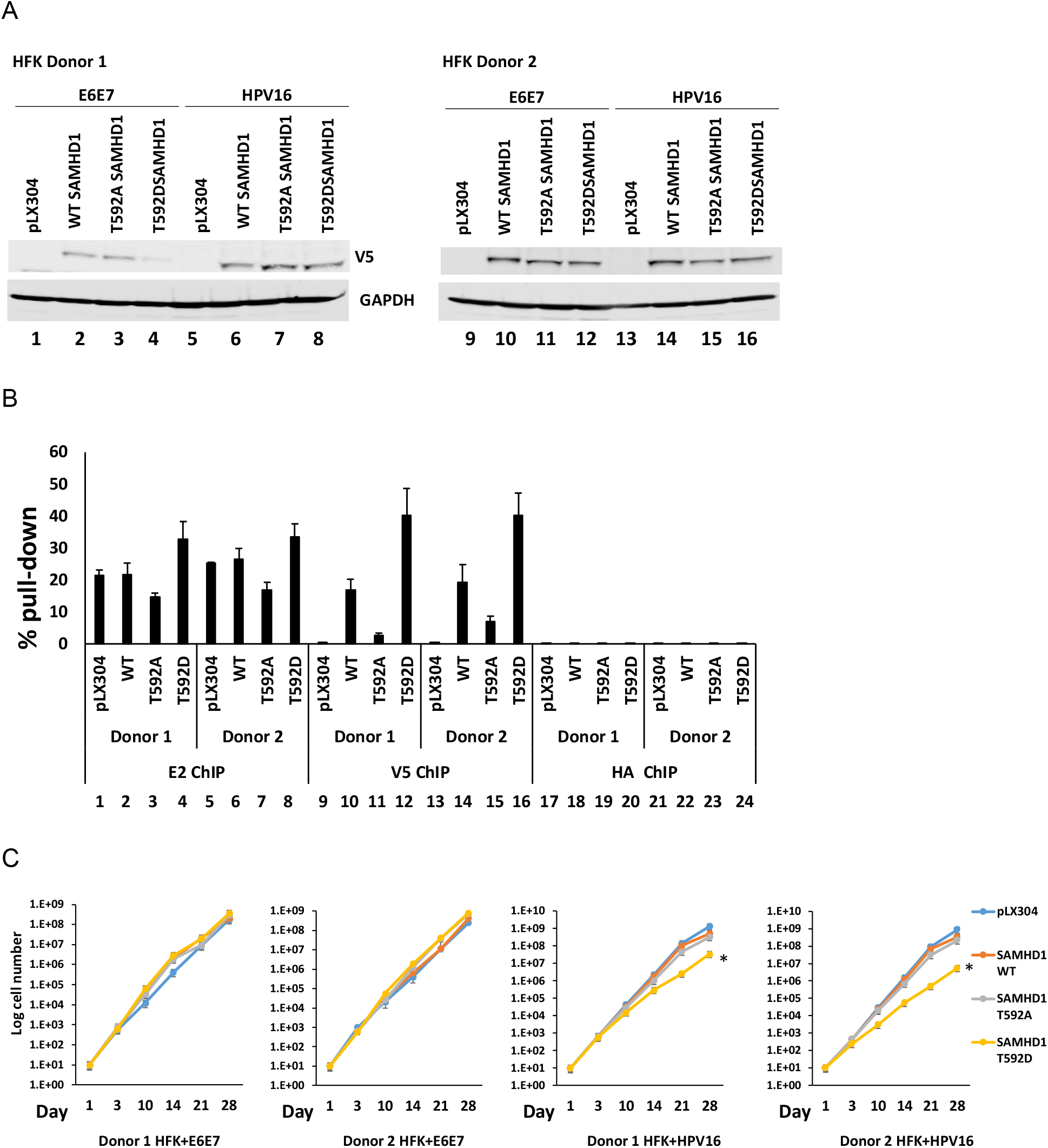
A. Western blot demonstrating expression of exogenous SAMHD1 proteins (detected with V5 antibody) in the indicated cell lines. B. Chromatin immunoprecipitation assays demonstrate that SAMHD1 T592D had increased recruitment to viral DNA than exogenous wild type SAMHD1 (p-value < 0.05). C. Growth curve of the indicated cell lines. Exogenous SAMHD1 T592D significantly (*, p-value <0.05) reduced the growth rate in HFK+HPV16 cell lines, but not in HFK+E6E7 cells.

Following differentiation of HFK+HPV16 cells, SAMHD1 T592D blocked the formation of viral replication foci (detected using γ-H2AX, Figure 4A). Growing, undifferentiated HFK+HPV16 cells had no γ-H2AX foci, HFK+E6E7 cells had no foci irrespective of differentiation status (not shown). Expression of SAMHD1 WT or SAMHD1 T592A had no effect on foci formation (Figure 4A), and quantitation of foci numbers confirmed only SAMHD1 T592D could block foci formation (Figure 4B). The quantitation (Figure 4B) summarizes the data from two independent HFK donor lines, donor 1 is shown in Figure 4A. The expression of the SAMHD1 proteins do not affect differentiation as involucrin induction was similar in all samples (Figure 4C).

**Figure 4.**
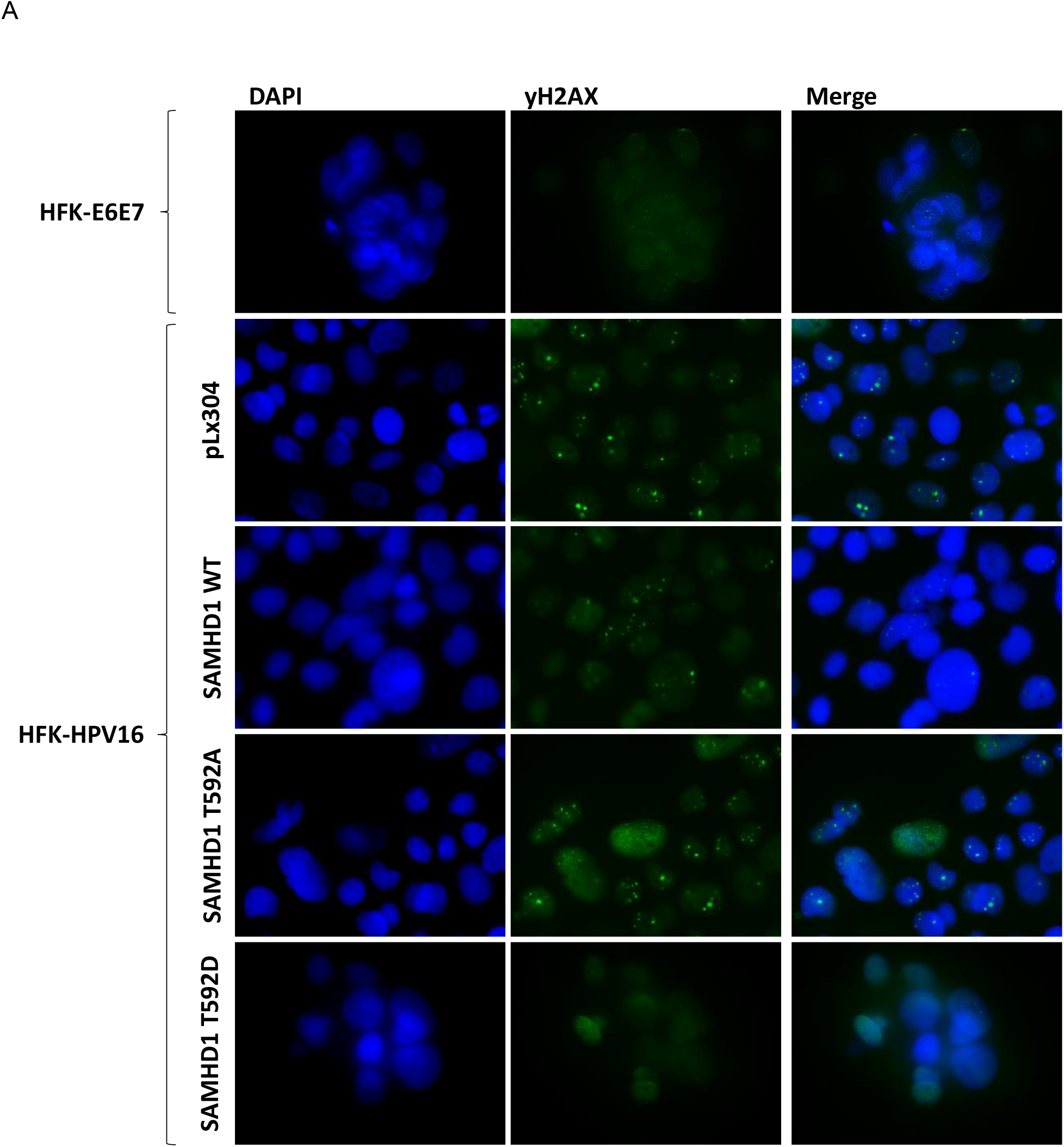
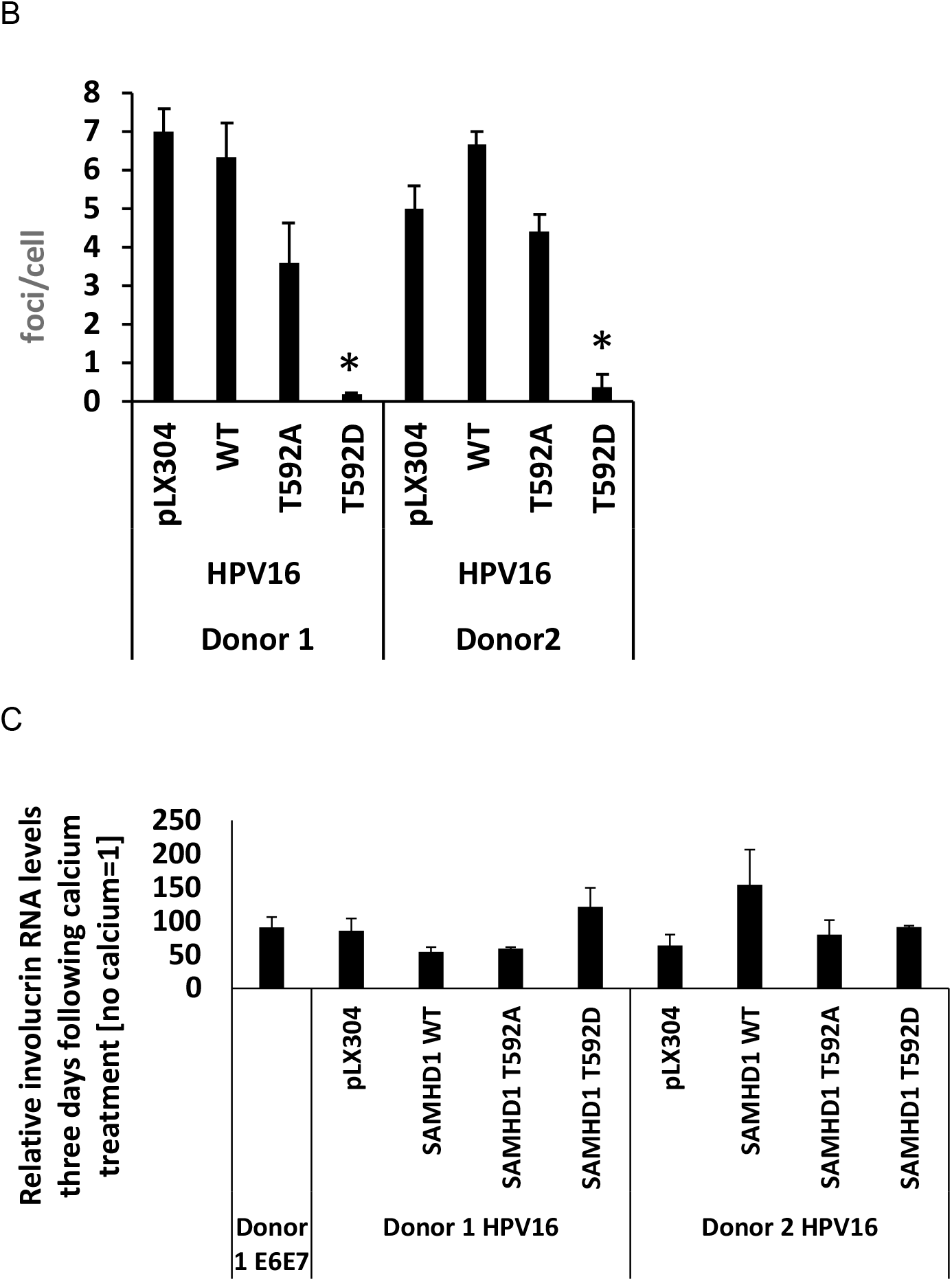
A. Calcium was added to the indicated cell lines to differentiate them and cells were fixed three days later and stained for γH2AX as a marker of viral replication foci. Donor 1 is shown, and Donor 2 cells behaved in a similar manner. B. The number of cells with γH2AX foci was quantitated and expressed as a percentage of all cells. There was a significant reduction in the percentage of cells with γH2AX foci in the SAMHD1 T592D samples (*, p-value < 0.05) when compared with all other samples. C. RNA was prepared from the differentiated cells and involucrin levels determined. The results are expressed relative to no calcium treatment = 1. In all cases there is a significant increase in involucrin RNA (p-value < 0.05), confirming that differentiation has not been affected by expression of any of the SAMHD1 proteins.

The results in Figures 1-4 demonstrate that SAMHD1 is recruited to E1-E2 replicating DNA, and that a T592D mutant has increased recruitment to the replicating DNA and can disrupt viral replication.

### Endothall, a protein phosphatase 2A (PP2A) inhibitor, promotes recruitment of endogenous SAMHD1 to replicating viral DNA

PP2A dephosphorylates SAMHD1 during transition from mitosis (37) and endothall is an inhibitor of PP2A (38). As SAMHD1 T592D (mimicking constant phosphorylation of SAMHD1) disrupts E1-E2 mediated DNA replication and viral genome amplification during mitosis, the ability of endothall to “mimic” SAMHD1 T592D by inducing increased endogenous SAMHD1 phosphorylation was tested. Figure 5A demonstrates that endothall significantly reduces the levels of E1-E2 DNA replication in transient C33a transfection experiments. Endothall did not induce death of the transfected cells as the transfected E1 plasmid levels were the same in the vehicle and endothall treated cells (Figure 5B). Endothall induced increased levels of E2, E1 and endogenous SAMHD1 recruited to the E1-E2 replicating DNA (Figure 5C). The attenuation of replication and the hyper-recruitment to the replicating DNA following endothall treatment is similar to the results generated with SAMHD1 T592D (Figure 2).

**Figure 5.**
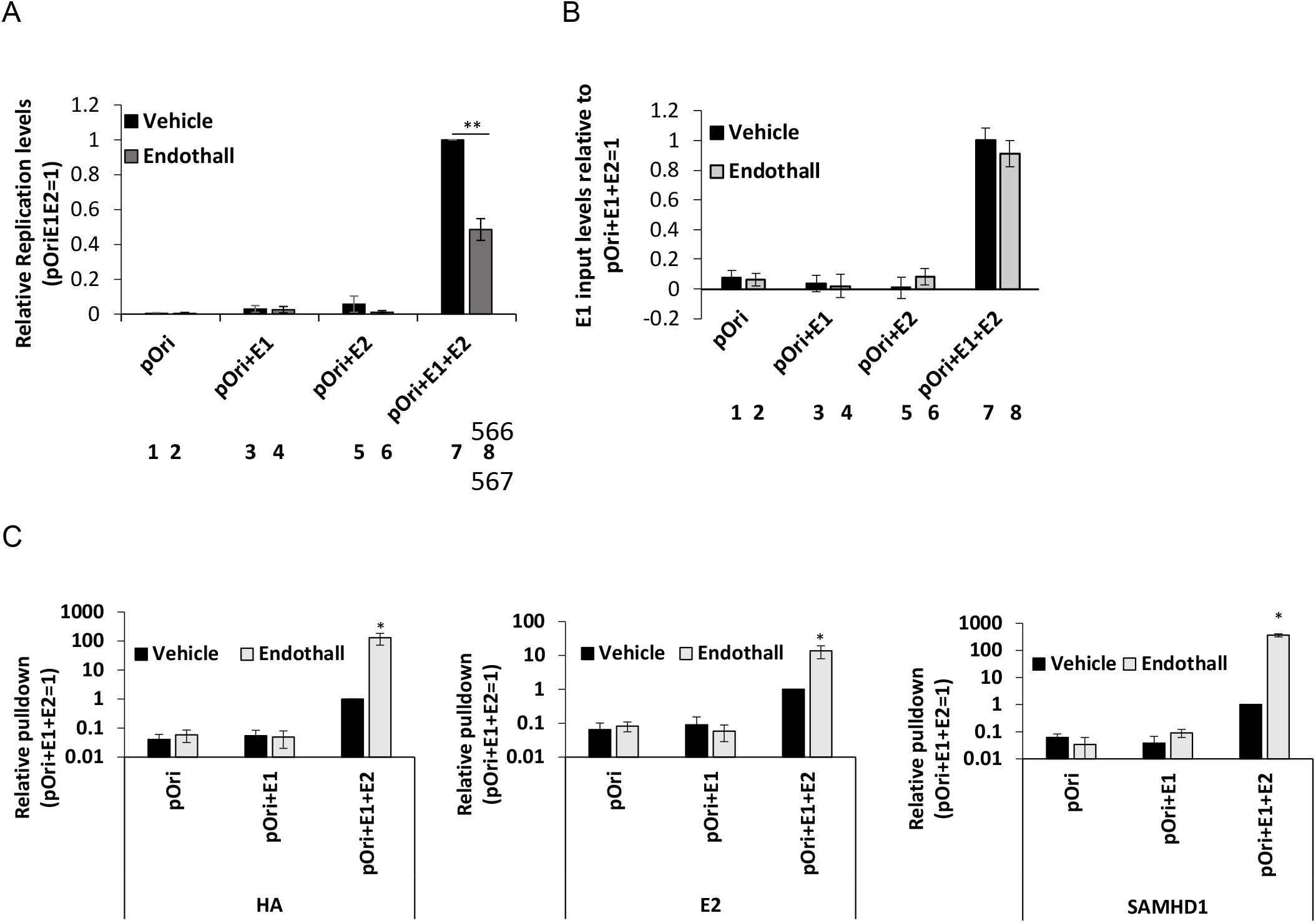
A. Replication assays were carried out in C33a cells transfected with the indicated plasmids; substantial replication is only detected in the presence of E1 and E2 expression (lanes 7 and 8). Treatment with endothall significantly reduced the replication signal (**, compare lane 8 with lane 7, p-value < 0.05). B. The levels of E1 input plasmid were determined in the control and endothall treated cells, demonstrating no reduction in E1 levels following drug treatment (lanes 7 and 8). This demonstrates that the combination of transfection and endothall treatment is not killing the transfected cells. C. Chromatin was prepared from C33a cells transfected with the indicated plasmids and ChIP assays carried out with HA (E1), E2 and endogenous SAMHD1 antibodies. The treatment with endothall significantly increased the recruitment of all three proteins to the replicating DNA (*, p-value < 0.05).

### Endothall promotes recruitment of SAMHD1 to the viral genome in HFK+HPV16 cells, attenuates their growth, and blocks viral replication foci formation during differentiation

As endothall is attenuating E1-E2 mediated DNA replication in C33a calls, and promoting recruitment of endogenous SAMHD1 onto the replicating DNA, the ability of endothall to disrupt the growth of HFK+HPV16 and HFK+E6E7 cells from two independent donors was determined. Figure 6A demonstrates that endothall significantly attenuates the growth of HFK+HPV16 cells (left panel). It also attenuates the growth of HFK+E6E7 cells but less than HFK+HPV16 (right panel). Chromatin was prepared from the treated HFK+HPV16 cells and the interaction of E2 and SAMHD1 with the replicating viral DNA determined using ChIP assays (Figure 7B). For E2 and SAMHD1 there was a significant increase in recruitment to the viral DNA with donor 1 (compare lanes 2 to 1 and 6 to 5, respectively). For donor 2 there was a significant increase in SAMHD1 recruitment (compare lane 8 with 7) and although there was an increase in E2 recruitment it did not reach significance as the p-value was greater than 0.05. HA was used as a control antibody (lanes 9-12) and there was significantly less viral DNA pulled down when compared with E2 and SAMHD1, and there was no significant difference in HA signal following endothall treatment. In Figure 6C we confirmed that the endothall treatment increased SAMHD1 phosphorylation in the two donor HFK+HPV16 cell lines (compare lanes 2 and 4 with 1 and 3), while there was very low SAMHD1 phosphorylation in the HFK+E6E7 cells that was not altered by endothall treatment (lanes 5-8).

**Figure 6.**
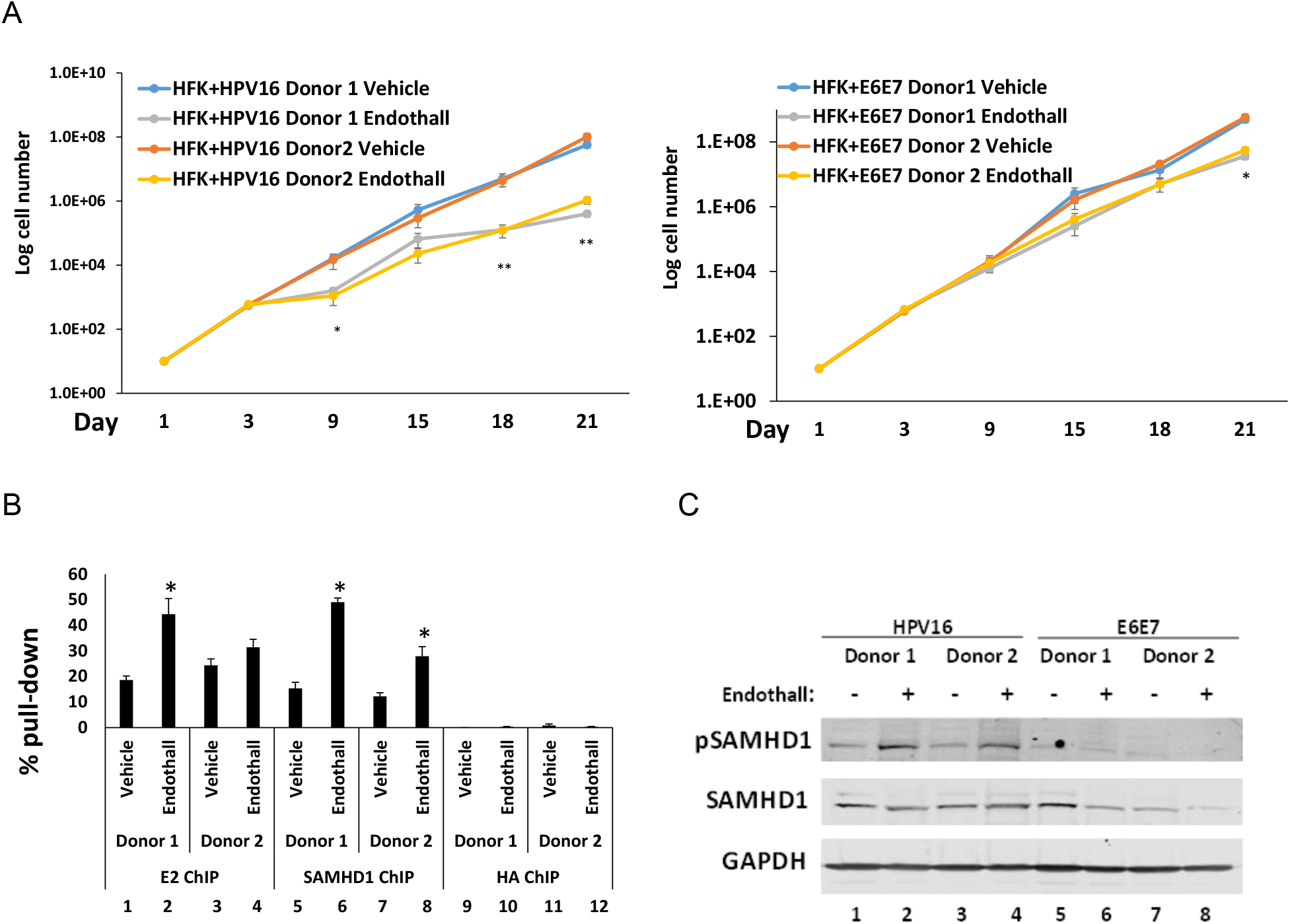
A. The indicated cell lines were treated with endothall (or vector control) over three weeks and cell growth monitored. In both HFK+HPV16 donors (left hand panel) there was a significant reduction in cellular growth at most time points tested in the endothall treated samples (*, p-value < 0.05). For HFK+E6E7 cells (right hand panel) there was only a significant reduction in cell growth at the 21 day time point with endothall treatment (*, p-value < 0.05). B. Chromatin was prepared from both HFK+HPV16 Donor lines and ChIP assays carried out with E2 and SAMHD1 (HA was used as a negative control). There is a significant increase in SAMHD1 recruitment to the viral DNA in the presence of endothall for both donors (*, p-value < 0.05). Donor 1 has a statistically significant increase in recruitment of E2 to the viral genome in the presence of endothall (*, p-value < 0.05). In Donor 2, there was an increase in E2 recruitment to the viral genome in the presence of endothall, but this did not reach significance. C. Treatment with endothall increased phosphorylation of SAMHD1 on T592 (pSAMHD1 signal) in both HFK+HPV16 donor lines, but did not do so in either HFK+E6E7 cell lines.

**Figure 7.**
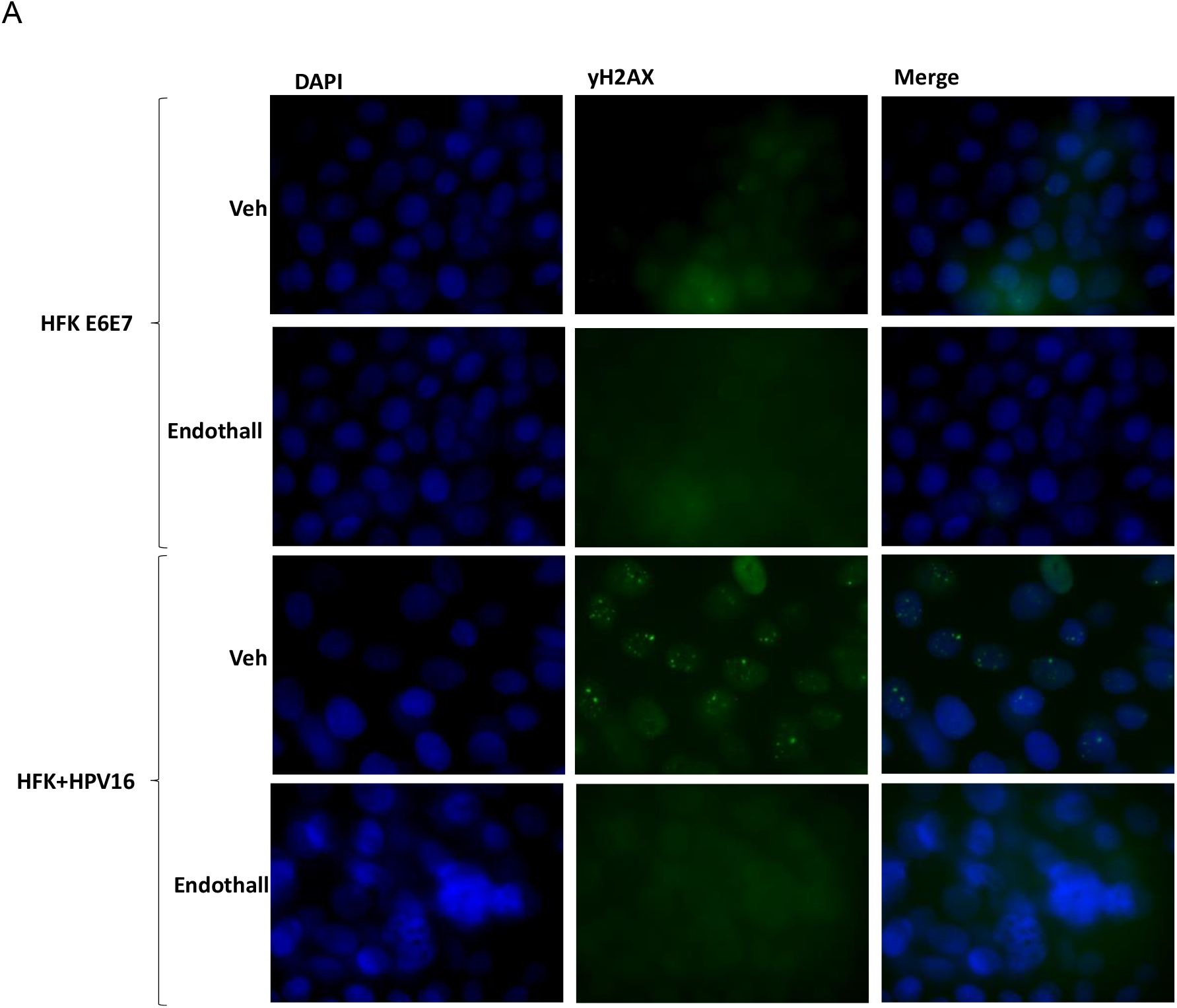
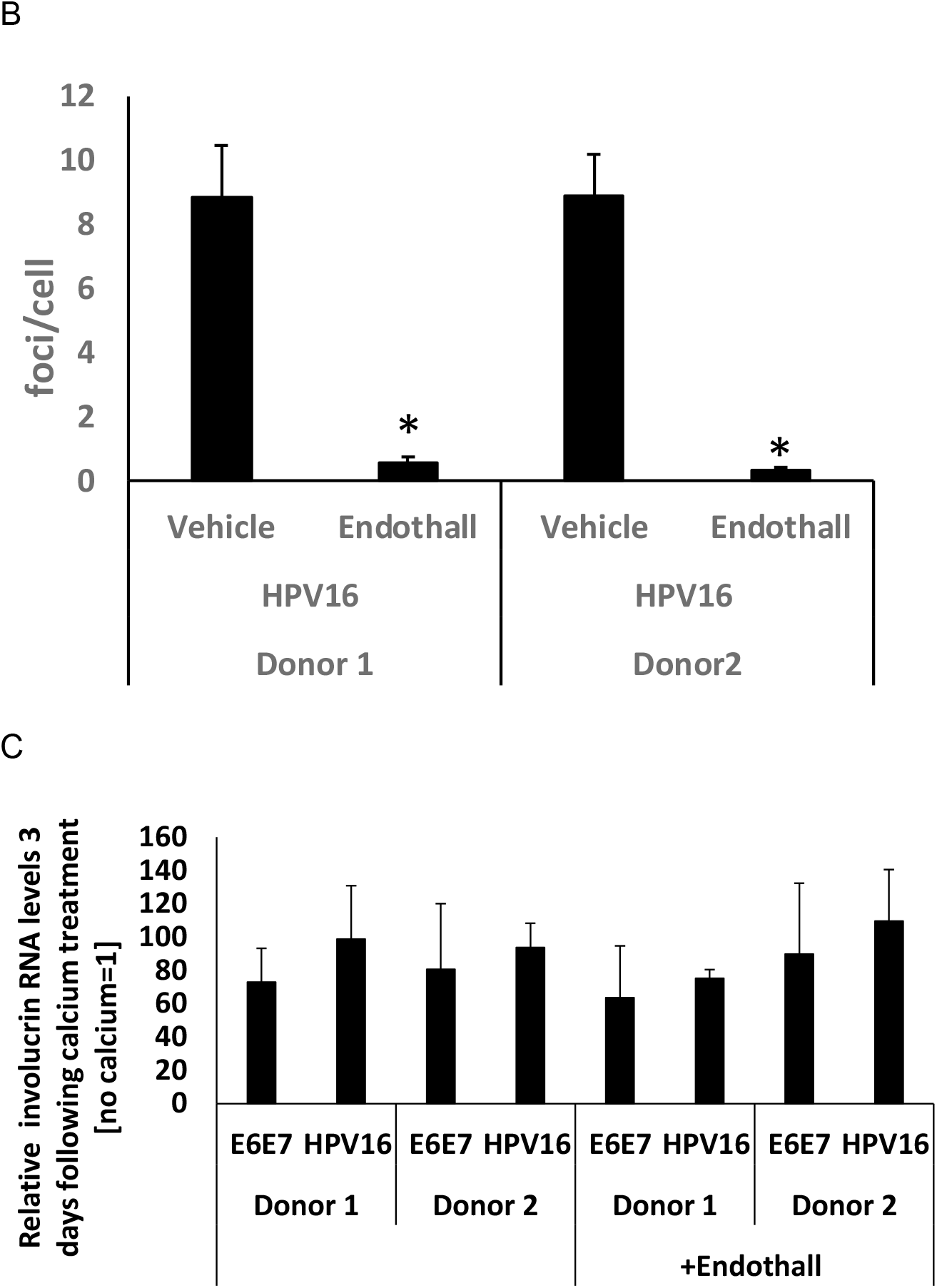
A. Calcium was added to the indicated cell lines to differentiate them in the presence of endothall or vehicle (Veh) and cells were fixed three days later and stained for γH2AX as a marker of viral replication foci. Donor 1 is shown, and donor 2 cells behaved in a similar manner. B. The number of cells with γH2AX foci was quantitated and expressed as a percentage of all cells. There was a significant reduction in the percentage of cells with γH2AX foci in the SAMHD1 T592D samples (*, p-value < 0.05) when compared with all other samples. C. RNA was prepared from the differentiated cells and involucrin levels determined. The results are expressed relative to no calcium treatment = 1. In all cases there is a significant increase in involucrin RNA (p-value < 0.05), confirming that differentiation has not been affected by endothall treatment.

The addition of endothall prevented replication foci formation following differentiation of HFK+HPV16 cells (Figure 7A). γH2AX is used as a marker for the viral replication foci, and there are no foci detectable in HFK+E6E7 cells (the top two sets of panels) irrespective of endothall treatment. As expected, in HFK+HPV16 replication foci developed three days following the addition of calcium (third panel set down). Endothall blocked formation of these replication foci (bottom panel set). The results shown are for donor 1 HFK cells, identical results were obtained in both donors and quantitation is shown in Figure 7B. Treatment with endothall did not affect differentiation, as involucrin induction was similar in all samples (Figure 7C).

### Endothall preferentially attenuates the growth of a HPV16 positive head and neck cancer cell line with episomal viral genomes

To determine whether the endothall growth attenuation results with HFK+HPV16 (Figure 6) could be extended to HPV16 positive head and neck cancer cell lines we treated three cell lines with endothall: HN30 which are HPV negative and p53 wild type; SCC-104 which are HPV16 positive and contain episomal viral genomes; and SCC-47 which are HPV16 positive and contain integrated viral genomes. The three cell lines were also treated with cisplatin. The IC50 for the treatments are shown in Figure 8A. All cell lines were equally sensitive to cisplatin while UMSCC104 was statistically more sensitive to endothall than the other two cell lines. Chromatin was prepared from the cells and the recruitment of SAMHD1 and E2 to the viral genome in UMSCC104 was determined with and without endothall treatment (Figure 8B). Endothall statistically increased the recruitment of E2 to the viral genome in UMSCC104 (right hand panel) and increased SAMHD1 recruitment, although it marginally failed to reach statistical significance (left had panel, p-value = 0.0505). The HA antibody was used as a control and resulted in background signals that were not varied by endothall.

**Figure 8.**
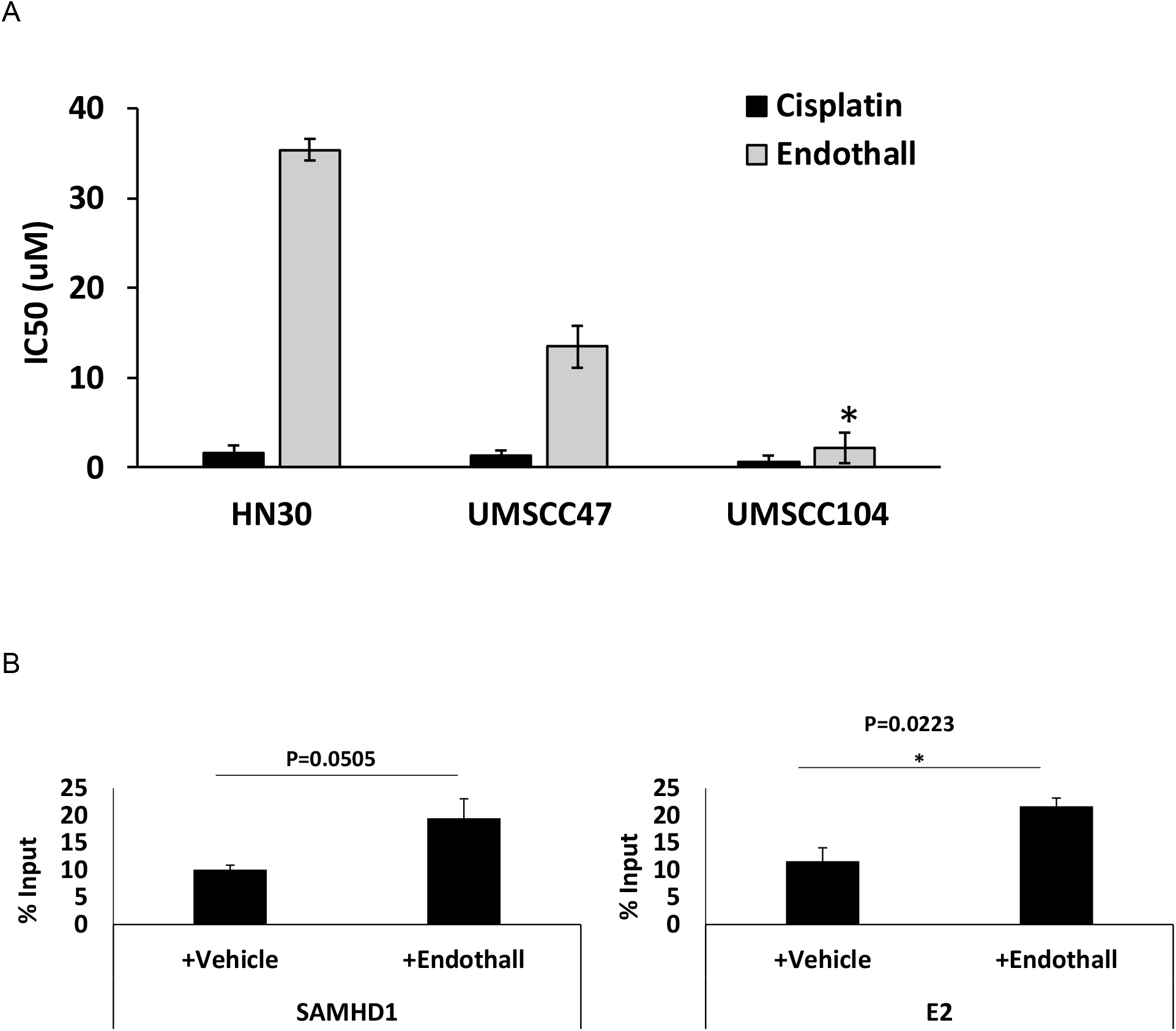
A. The indicated cells were treated with endothall and cisplatin and the IC50 determined. All three cell lines were equally sensitive to cisplatin, while UMSCC104 (which contains episomal HPV16 genomes) was significantly more sensitive to endothall than the other two lines (*, p-value < 0.05). B. Chromatin was prepared from UMSC104 cells treated with and without endothall and ChIP assays carried out for SAMHD1 (left panel) and E2 (right panel). E2 recruitment was significantly increased in the presence of endothall (*, p-value < 0.05), and while SAMHD1 recruitment was also increased it failed to reach significance.

## Discussion

HPV activates the DNA damage response (DDR) to promote the viral life cycle, and inhibition of the DDR can block viral genome amplification (16). One reason the virus activates the DDR is to promote HR replication of the viral genome, allowing amplification in the presence of an active DDR (22). A host of HR factors are recruited to the replicating viral genome and are critical for the viral life cycle (7, 8, 10, 12–14, 19, 44). Here we demonstrate that viral replication converts SAMHD1 into an HR factor via phosphorylation on threonine 592, the viral oncogenes E6 and E7 are unable to do this. It may not necessarily by viral replication *per se* that converts SAMHD1 into an HR factor but expression of the viral helicase E1, a known activator of the DDR when overexpressed (45-47). The role of SAMHD1 in HR is to promote DNA end resection that generates substrates required for HR via recruitment of CtIP protein to sites of DNA damage (35). In addition to regulation by phosphorylation, the role of SAMHD1 in HR can be controlled by SIRT1 deacetylation as this facilitates SAMHD1 binding to double strand DNA breaks (36). Moreover, SAMHD1 SUMOylation by PIAS1 also regulates its DNA binding and anti-viral activity (48). SIRT1 is a known component of the HPV16 E1-E2 DNA replication complex that can deacetylate E2 and HR host factors such as WRN to promote high fidelity E1- E2 DNA replication and control the viral life cycle (7-9, 49). Overall, the results suggest that HPV16 replication (or replication factors) convert SAMHD1 into an HR factor that assists with replication of the viral genome. We demonstrate here that SAMHD1 is recruited to E1-E2 replicating DNA in C33a cells, and also to the HPV16 genome in HFK cells immortalized by HPV16. CRISPR/Cas9 removal of SAMHD1 in N/Tert-1 cells containing the HPV16 genome resulted in increased cell proliferation, viral replication and DNA damage demonstrating that SAMHD1 is a restriction factor controlling the virally infected cell (33). The phenotypes observed could also be related to the effects on host DNA replication in the absence of SAMHD1. If the virus converts SAMHD1 into an HR factor this could be important in managing host DNA replication in the presence of an active DDR generated by viral replication.

While HPV genomes integrate in the majority of cervical cancers, in oropharyngeal cancers the majority retain an episomal HPV16 genome (50, 51). The presence of viral replication in these cancers presents a potential therapeutic target as reducing viral replication will ultimately reduce the levels of E6 and E7 proteins potentially increasing the expression of their tumor suppressor targets p53 and pRb, respectively. Such an increase would promote the response of the tumors to chemotherapeutic agents such as radiation and cisplatin. The conversion of SAMHD1 into an HR factor only in the HFK cells containing replicating viral genomes creates a difference with control cells that we sought to exploit. Phosphatases are an important component of DNA replication complexes and we investigated whether a SAMHD1 phosphor-mimetic could disrupt HPV16 replication. The results demonstrate that SAMHD1 T592D (the aspartic acid providing a negative charge mimicking phosphorylation) is hyper-recruited to E1-E2 replicating DNA in C33a cells, and reduces the levels of replication. This results supports the idea that dephosphorylation of SAMHD1 in the replication complex may cycle it off the replicating DNA, allowing continuous replication. SAMHD1 T592D also promoted recruitment of E1 and E2 to the replicating DNA in C33a cells suggesting that the replication complex has become stalled, as there is a significant reduction in replication even though more replication factors are being recruited to the replicating DNA. In HFK+HPV16 cells, SAMHD T592D reduced the growth rates of the cells but could not do this in HFK+E6E7 cells. SAMHD1 T592D is also hyper-recruited to the viral genome in HFK cells and promotes an increase in the recruitment of E2 to the viral genome. The presence of SAMHD1 T592D also prevented the formation of viral replication centers following differentiation of HFK+HPV16 cells, as demonstrated by a significant reduction in γH2AX foci development in these cells following calcium treatment when compared with control cells. This demonstrates that SAMHD1 T592D can disrupt viral replication during differentiation. Although the HFK+HPV16 SAMHD1 T592D cells grew slower we did not see a significant reduction in the viral DNA in these cells. This is perhaps due to cells dying due to reduced viral DNA over the longer term of these growth assays, or it could be due to SAMHD1 T592D regulating host DNA replication in the HFK+HPV16 cells and attenuating growth via slowing down host replication. It could be a combination of several factors that we will investigate in future studies.

To progress our results in a therapeutic direction we next investigated the ability of the phosphatase PP2A inhibitor endothall to attenuate HPV16 replication and cell growth. PP2A dephosphorylates SAMHD1 (37) and therefore we anticipated that PP2A inhibition would mimic the phenotypes of SAMHD T592D. This is indeed what we saw as endothall treatment increased SAMHD1, E1 and E2 recruitment to replicating DNA in our C33a model whilst attenuating replication levels; endothall attenuated the growth of HFK+HPV16 cells preferentially when compared with control cells; endothall increased the recruitment of SAMHD1 and E2 to viral DNA in HFK+HPV16 cells, and prevented the development of replication foci following calcium induced differentiation. Finally we demonstrate that growth of UMSCC104 cells, an HPV16 positive head and neck cancer cell line containing episomal viral genomes, is attenuated more than other head and neck cancer cell lines following endothall treatment, and SAMHD1 and E2 recruitment to the viral genome is increased by endothall treatment of UMSCC104 cells.

Overall our results suggest that phosphatase inhibitors are therapeutic candidates for the management of HPV16 positive cancers that contain episomal genomes. While we have demonstrated that endothall treatment increases the recruitment of SAMHD1 to E1-E2 replicating DNA, endothall treatment also likely increases the levels of recruitment of additional DDR/HR factors to the viral DNA. In addition, the toxicity of endothall in the HPV16 full genome cells could be related to the targeting of host replication, which is operating in an environment of an active DDR that would ordinarily arrest DNA replication. Future work will continue investigating the potential of targeting phosphatases for the treatment of HPV malignancies that contain replicating DNA.

## Materials and methods

### Cell culture, plasmids, and reagents

N/Tert-1 cells were cultured in keratinocyte serum-free medium (K-SFM) (Invitrogen; catalog no. 37010022) supplemented with bovine pituitary extract, EGF (Invitrogen), 0.3 mM calcium chloride (MilliporeSigma; 21115) and 7.5 uM hygromycin at 37 °C in a 5% CO_2_/95% air atmosphere. Human foreskin keratinocytes (HFKs) were immortalized with HPV16 or with 16 E6E7 expression vector pLXSN16E6E7, a gift from Denise Galloway (Addgene plasmid #52394), as described previously (21). HFKs were cultured in DermaLife-K Complete media (LifeLine Cell Technologies). Mitomycin C-treated 3T3-J2 fibroblasts feeders were plated 24 hours prior to seeding N/Tert-1 or HFK cells on top of the feeders, in their respective cell culture media. Media was refreshed and 3T3-J2s supplemented as required. HN30 and UMSCC47 cells were cultured in DMEM supplemented with 10% FBS (R&D Systems). UMSCC104 were cultured in EMEM (Quality Biological Inc) supplemented with 20% FBS and NEAA (ThermoFisher). Both UMSCC104 and UMSCC47 cell lines were obtained from Millipore. HN30 cells were a gift from Hisashi Harada. C33a cells were obtained from ATCC (Manassas, VA, USA) and grown in Dulbecco’s Modified Eagle’s Medium (Invitrogen, Carlsbad, CA, USA) supplemented with 10% fetal bovine serum and were passaged every 3–4 days. In all cases, cell identity was confirmed via “fingerprinting”, and cell cultures were routinely monitored for mycoplasma.

To measure proliferation, cells were seeded in triplicate into 6 well plates at a density of 1×10^5^ cells per well, and grown to 80% confluency (typically 3 days). Cells were then harvested by trypsinization, stained with trypan blue and viable cells counted. 1×10^5^ cells per dish were re-plated and this was repeated every three-four days for 3 weeks. To measure the effect of endothall on proliferation, cells were cultured in either 5 μM endothall or vehicle in media throughout.

To measure sensitivity of cells to endothall, cells were seeded into 6 well plates at a density of 1×10^4^ cells per dish, and cultured in either vehicle, 1, 5, 10, 25, or 50uM endothall. Once vehicle wells reached 70-80% confluency, media was removed and cells were washed twice with PBS. 1 mL of 0.5% crystal violet was added per well, and plates incubated shaking at room temperature for 30 mins. Wells were washed five times with PBS and plates left to dry. Crystal violet images were scanned using the Odyssey^®^ CLx Imaging System and ImageJ was used for quantification.

All of the HPV16 plasmids utilized in these studies have been previously used and described by this laboratory: HPV16 pOriLacZ (pOriLacZ), HPV16 E1-(hemagglutinin, HA) (E1), HPV16 E2 (10, 39). SAMHD1 WT, T592A and T592D lentiviral plasmids were described previously (32)

### Transient DNA Replication Assay

C33a cells were plated out at 5×10^5^in 10 cm dishes. The following day, plasmid DNA was transfected using the calcium phosphate method. Three days post-transfection, low molecular weight DNA was extracted using the Hirt method as previously described by Boner *at al*. 2002. The digested sample was extracted twice with phenol:chloroform:isoamyl alcohol (25:24:1) and precipitated with ethanol. Following centrifugation, the DNA pellet was washed with 70% ethanol, dried and resuspended in a total of 150 µL water. Forty two microliters of sample were digested with DpnI (New England Biolabs, Ipswich, MA, USA) overnight to remove unreplicated pOri16LacZ; the sample was then digested with ExoIII (New England Biolabs) for 1 hr. Replication was determined by real-time PCR, as described previously (39). For endothall treatment, cells were cultured in 5uM endothall 24 hours post-transfection, which was also 48 hours before harvesting.

### Western blotting

Specified cells were trypsinized, washed with PBS and resuspended in 2x pellet volume NP40 protein lysis buffer (0.5% Nonidet P-40, 50 mM Tris [pH 7.8], 150 mM NaCl) supplemented with protease inhibitor (Roche Molecular Biochemicals) and phosphatase inhibitor cocktail (MilliporeSigma). Cell suspension was incubated on ice for 20 min and then centrifuged for 20 min at 184,000 rcf at 4 °C. Protein concentration was determined using the Bio-Rad protein estimation assay according to manufacturer’s instructions. 50 μg protein was mixed with 2x Laemmli sample buffer (Bio-Rad) and heated at 95 °C for 5 min. Protein samples were separated on <u>Novex</u> 4–12% Tris-glycine gel (Invitrogen) and transferred onto a nitrocellulose membrane (Bio-Rad) at 30V overnight using the wet-blot transfer method. Membranes were then blocked with Odyssey (PBS) blocking buffer (diluted 1:1 with PBS) at room temperature for 1 hr and probed with indicated primary antibody diluted in Odyssey blocking buffer, overnight. Membranes were washed with PBS supplemented with 0.1% Tween (PBS-Tween) and probed with the Odyssey secondary antibody (goat anti-mouse IRdye 800CW or goat anti-rabbit IRdye 680CW) (Licor) diluted in Odyssey blocking buffer at 1:10,000. Membranes were washed twice with PBS-Tween and an additional wash with 1X PBS. After the washes, the membrane was imaged using the Odyssey^®^ CLx Imaging System and ImageJ was used for quantification, utilizing GAPDH as internal loading control. Primary antibodies used for western blotting studies are as follows: 16E2 monoclonal B9 1/500 (52), SAMHD1 1/1000 (Cell Signaling Technology, 49158), phospho-SAMHD1 1/500 (Cell Signaling Technology, 89930) GAPDH 1/10,000 (Santa Cruz, sc-47724), HA-Tag (Abcam, ab9110), WRN 1/1000 (Cell Signaling Technology, 4666), MRE11 1/500 (Cell Signaling Technology, 4847), V5-Tag 1/1000 (Life Technologies, A190- 119A).

### Immunoprecipitation assay

Cell lysate was prepared as described above. 250 µg of the lysate was incubated with lysis buffer (0.5% Nonidet P-40, 50mM Tris [pH 7.8], and 150mM NaCl), supplemented with protease inhibitor (Roche Molecular Biochemicals) and phosphatase inhibitor cocktail (MilliporeSigma) to a total volume of 500 µl. Primary antibody of interest or a HA-tag antibody (used as a negative control) was added to this prepared lysate and rotated at 4°C overnight. The following day, 40 µl of protein A beads per sample (MilliporeSigma; equilibrated to lysis buffer as mentioned in the manufacturer’s protocol) was added to the above mixture and rotated for another 4 hours at 4°C. The samples were gently washed with 500 µl lysis buffer by centrifugation at 1000 x g for 2-3 min. This wash was repeated 3 times. The bead pellet was resuspended in 4X Laemmli sample buffer (Bio-Rad), heat denatured and centrifuged at 1000 *rcf* for 2-3 min. The supernatant applied to an SDS-PAGE system to separate and resolve proteins, and was then transferred onto a nitrocellulose membrane using wet-blot transfer method. The membrane was probed for the presence of specific proteins as mentioned in the description of western blotting above.

### Chromatin immunoprecipitation (ChIP)

Cross-linking and chromatin extraction of C33a cells was carried out as previously described (10). For chromatin extraction from keratinocytes, the ChIP-It Enzymatic kit (Active Motif) was utilized, as the protocol dictated, including dounce homogenisation to disrupt cells and incubation with shearing enzymes for 7.5 minutes with frequent agitation. In both cases, sheared chromatin was incubated with 1 ug primary antibody (HA, Abcam ab9110; V5 Abcam ab15828; SAMHD1, ProteinTech 1258-6-AP; a sheep E2 antibody as previously described (10)) and protein-G conjugated magnetic beads, rotating overnight at 4°C. Beads were then washed and DNA eluted via Proteinase K digestion. Precipitated DNA was measured via qPCR (Fwd 5’- GAAAACGAAAAGCTACACCCA-3’, Rev 5’-CAATGAATAACCACAACACAATTA-3’), and percentage pulldown calculated by calculation to input:% *Pulldown* = 100 × 2^*Input Ct−ChIP Ct*^. For endothall treatment, cells were cultured in 5 uM endothall for 24 hours before fixation and chromatin harvest.

### Immunofluorescence

For fluorescent staining, cells were cultured on coverslips coated with poly-L-lysine (MilliporeSigma), and differentiated by culturing in 1.5 mM calcium-containing media (MCF-154, supplemented with CaCl_2_, both Invitrogen) for three days. For endothall treatment, cells were cultured in 5 uM endothall for the three days of differentiation. Coverslips were then washed in PBS and fixed by incubation with ice-cold methanol for 10 mins. These were permeabilized by incubation in 0.2% Triton X-100/PBS at room temperature for 15 minutes. Background staining was reduced by blocking coverslips in 10% Normal goat serum (LifeTechnologies). Primary antibodies were used at the dilutions as follows: SAMHD1 (1/500, Cell Signaling Technology 49158), phospho-S139 Histone yH2AX 1/200 (Cell Signaling Technology, 9718). Immune complexes were visualized using Alexa 488- or Alexa 595-labeled anti-species specific antibody conjugates (Molecular Probes). Cellular DNA was stained with 4’,6-diamidino-2-phenylindole (DAPI, Santa Cruz sc-3598). Coverslips were mounted onto slides using Vectashield medium (ThermoFisher) and visualized, and fluorescence quantified using the Keyence imaging system.

Differentiation was confirmed by qRTPCR analysis for involucrin (Fwd 5’- TCCTCCAGTCAATACCCATCAG-3’, Rev 5’- CAGCAGTCATGTGCTTTTCCT-3’). Involucrin induction was calculated relative to undifferentiated control, using GAPDH (Fwd 5’- GGAGCGAGATCCCTCCAAAAT-3’, Rev 5’- GGCTGTTGTCATACTTCTCATGG-3’) as internal housekeeping control.

### Statistics

Standard error was calculated from three independent experiments and significance determined using a student’s t-test.

## Acknowledgements

This work was supported by US NIH grant R01DE029471 (IMM) and R01AI141410 (RL). RL was supported by Research Scholar Grant (134703-RSG-20–054-01-MPC) and Institutional Research Grant (IRG-14-192-40) from the American Cancer Society. The funders had no role in study design, data collection and analysis, decision to publish, or preparation of the manuscript.

## References

1. zur Hausen H. 2009. Papillomaviruses in the causation of human cancers - a brief historical account. Virology 384:260–265.

2. McBride AA. 2013. The Papillomavirus E2 proteins. Virology 445:57–79.

3. Bergvall M, Melendy T, Archambault J. 2013. The E1 proteins. Virology 445:35–56.

4. Park P, Copeland W, Yang L, Wang T, Botchan MR, Mohr IJ. 1994. The cellular DNA polymerase alpha-primase is required for papillomavirus DNA replication and associates with the viral E1 helicase. Proceedings of the National Academy of Sciences of the United States of America 91:8700–8704.

5. Clower RV, Fisk JC, Melendy T. 2006. Papillomavirus E1 protein binds to and stimulates human topoisomerase I. Journal of virology 80:1584–1587.

6. Masterson PJ, Stanley MA, Lewis AP, Romanos MA. 1998. A C-terminal helicase domain of the human papillomavirus E1 protein binds E2 and the DNA polymerase alpha-primase p68 subunit. Journal of virology 72:7407–7419.

7. Das D, Bristol ML, Smith NW, James CD, Wang X, Pichierri P, Morgan IM. 2019. Werner Helicase Control of Human Papillomavirus 16 E1-E2 DNA Replication Is Regulated by SIRT1 Deacetylation. MBio 10.

8. Das D, Smith N, Wang X, Morgan IM. 2017. The Deacetylase SIRT1 Regulates the Replication Properties of Human Papillomavirus 16 E1 and E2. Journal of virology 91:e00102–e00117.

9. James CD, Das D, Morgan EL, Otoa R, Macdonald A, Morgan IM. 2020. Werner Syndrome Protein (WRN) Regulates Cell Proliferation and the Human Papillomavirus 16 Life Cycle during Epithelial Differentiation. mSphere 5.

10. Gauson EJ, Donaldson MM, Dornan ES, Wang X, Bristol M, Bodily JM, Morgan IM. 2015. Evidence supporting a role for TopBP1 and Brd4 in the initiation but not continuation of human papillomavirus 16 E1/E2 mediated DNA replication. Journal of virology 89:17684–17699.

11. Donaldson MM, Mackintosh LJ, Bodily JM, Dornan ES, Laimins LA, Morgan IM. 2012. An interaction between human papillomavirus 16 E2 and TopBP1 is required for optimum viral DNA replication and episomal genome establishment. Journal of virology 86:12806–12815.

12. Chappell WH, Gautam D, Ok ST, Johnson BA, Anacker DC, Moody CA. 2015. Homologous Recombination Repair Factors, Rad51 and BRCA1, are Necessary for Productive Replication of Human Papillomavirus 31. Journal of virology doi:JVI.02495-15 [pii].

13. Anacker DC, Gautam D, Gillespie KA, Chappell WH, Moody CA. 2014. Productive replication of human papillomavirus 31 requires DNA repair factor Nbs1. Journal of virology 88:8528–8544.

14. Gillespie Ka, Mehta KP, Laimins La, Moody Ca. 2012. Human papillomaviruses recruit cellular DNA repair and homologous recombination factors to viral replication centers. Journal of virology 86:9520–6.

15. James CD, Fontan CT, Otoa R, Das D, Prabhakar AT, Wang X, Bristol ML, Morgan IM. 2020. Human Papillomavirus 16 E6 and E7 Synergistically Repress Innate Immune Gene Transcription. mSphere 5.

16. Moody CA, Laimins LA. 2009. Human papillomaviruses activate the ATM DNA damage pathway for viral genome amplification upon differentiation. PLoS pathogens 5:e1000605.

17. Anacker DC, Moody CA. 2017. Modulation of the DNA damage response during the life cycle of human papillomaviruses. Virus research 231:41–49.

18. Gautam D, Moody CA. 2016. Impact of the DNA Damage Response on Human Papillomavirus Chromatin. PLoS pathogens 12:e1005613.

19. Prabhakar AT, James CD, Fontan CT, Otoa R, Wang X, Bristol ML, Hill RD, Dubey A, Morgan IM. 2023. Human Papillomavirus 16 E2 Interaction with TopBP1 Is Required for E2 and Viral Genome Stability during the Viral Life Cycle. J Virol 97:e0006323.

20. Prabhakar AT, James CD, Das D, Fontan CT, Otoa R, Wang X, Bristol ML, Morgan IM. 2022. Interaction with TopBP1 Is Required for Human Papillomavirus 16 E2 Plasmid Segregation/Retention Function during Mitosis. J Virol 96:e0083022.

21. Prabhakar AT, James CD, Das D, Otoa R, Day M, Burgner J, Fontan CT, Wang X, Glass SH, Wieland A, Donaldson MM, Bristol ML, Li R, Oliver AW, Pearl LH, Smith BO, Morgan IM. 2021. CK2 Phosphorylation of Human Papillomavirus 16 E2 on Serine 23 Promotes Interaction with TopBP1 and Is Critical for E2 Interaction with Mitotic Chromatin and the Viral Life Cycle. mBio doi:10.1128/mBio.01163-21:e0116321.

22. Sakakibara N, Chen D, McBride AA. 2013. Papillomaviruses use recombination-dependent replication to vegetatively amplify their genomes in differentiated cells. PLoS pathogens 9:e1003321.

23. Goldstone DC, Ennis-Adeniran V, Hedden JJ, Groom HC, Rice GI, Christodoulou E, Walker PA, Kelly G, Haire LF, Yap MW, de Carvalho LP, Stoye JP, Crow YJ, Taylor IA, Webb M. 2011. HIV-1 restriction factor SAMHD1 is a deoxynucleoside triphosphate triphosphohydrolase. Nature 480:379–82.

24. Kim CA, Bowie JU. 2003. SAM domains: uniform structure, diversity of function. Trends Biochem Sci 28:625–8.

25. Beloglazova N, Flick R, Tchigvintsev A, Brown G, Popovic A, Nocek B, Yakunin AF. 2013. Nuclease activity of the human SAMHD1 protein implicated in the Aicardi-Goutieres syndrome and HIV-1 restriction. J Biol Chem 288:8101–10.

26. Ji X, Wu Y, Yan J, Mehrens J, Yang H, DeLucia M, Hao C, Gronenborn AM, Skowronski J, Ahn J, Xiong Y. 2013. Mechanism of allosteric activation of SAMHD1 by dGTP. Nat Struct Mol Biol 20:1304–9.

27. Lahouassa H, Daddacha W, Hofmann H, Ayinde D, Logue EC, Dragin L, Bloch N, Maudet C, Bertrand M, Gramberg T, Pancino G, Priet S, Canard B, Laguette N, Benkirane M, Transy C, Landau NR, Kim B, Margottin-Goguet F. 2012. SAMHD1 restricts the replication of human immunodeficiency virus type 1 by depleting the intracellular pool of deoxynucleoside triphosphates. Nat Immunol 13:223–228.

28. Ryoo J, Choi J, Oh C, Kim S, Seo M, Kim SY, Seo D, Kim J, White TE, Brandariz-Nunez A, Diaz-Griffero F, Yun CH, Hollenbaugh JA, Kim B, Baek D, Ahn K. 2014. The ribonuclease activity of SAMHD1 is required for HIV-1 restriction. Nat Med 20:936–41.

29. St Gelais C, de Silva S, Amie SM, Coleman CM, Hoy H, Hollenbaugh JA, Kim B, Wu L. 2012. SAMHD1 restricts HIV-1 infection in dendritic cells (DCs) by dNTP depletion, but its expression in DCs and primary CD4+ T-lymphocytes cannot be upregulated by interferons. Retrovirology 9:105.

30. Kim ET, Roche KL, Kulej K, Spruce LA, Seeholzer SH, Coen DM, Diaz-Griffero F, Murphy EA, Weitzman MD. 2019. SAMHD1 Modulates Early Steps during Human Cytomegalovirus Infection by Limiting NF-kappaB Activation. Cell Rep 28:434–448.e6.

31. Laguette N, Sobhian B, Casartelli N, Ringeard M, Chable-Bessia C, Segeral E, Yatim A, Emiliani S, Schwartz O, Benkirane M. 2011. SAMHD1 is the dendritic-and myeloid-cell-specific HIV-1 restriction factor counteracted by Vpx. Nature 474:654–7.

32. Zhang K, Lv DW, Li R. 2019. Conserved Herpesvirus Protein Kinases Target SAMHD1 to Facilitate Virus Replication. Cell Rep 28:449–459.e5.

33. James CD, Prabhakar AT, Otoa R, Evans MR, Wang X, Bristol ML, Zhang K, Li R, Morgan IM. 2019. SAMHD1 Regulates Human Papillomavirus 16-Induced Cell Proliferation and Viral Replication during Differentiation of Keratinocytes. mSphere 4.

34. Cabello-Lobato MJ, Wang S, Schmidt CK. 2017. SAMHD1 Sheds Moonlight on DNA Double-Strand Break Repair. Trends Genet 33:895–897.

35. Daddacha W, Koyen AE, Bastien AJ, Head PE, Dhere VR, Nabeta GN, Connolly EC, Werner E, Madden MZ, Daly MB, Minten EV, Whelan DR, Schlafstein AJ, Zhang H, Anand R, Doronio C, Withers AE, Shepard C, Sundaram RK, Deng X, Dynan WS, Wang Y, Bindra RS, Cejka P, Rothenberg E, Doetsch PW, Kim B, Yu DS. 2017. SAMHD1 Promotes DNA End Resection to Facilitate DNA Repair by Homologous Recombination. Cell Rep 20:1921–1935.

36. Kapoor-Vazirani P, Rath SK, Liu X, Shu Z, Bowen NE, Chen Y, Haji-Seyed-Javadi R, Daddacha W, Minten EV, Danelia D, Farchi D, Duong DM, Seyfried NT, Deng X, Ortlund EA, Kim B, Yu DS. 2022. SAMHD1 deacetylation by SIRT1 promotes DNA end resection by facilitating DNA binding at double-strand breaks. Nat Commun 13:6707.

37. Schott K, Fuchs NV, Derua R, Mahboubi B, Schnellbächer E, Seifried J, Tondera C, Schmitz H, Shepard C, Brandariz-Nuñez A, Diaz-Griffero F, Reuter A, Kim B, Janssens V, König R. 2018. Dephosphorylation of the HIV-1 restriction factor SAMHD1 is mediated by PP2A-B55α holoenzymes during mitotic exit. Nat Commun 9:2227.

38. Erdödi F, Tóth B, Hirano K, Hirano M, Hartshorne DJ, Gergely P. 1995. Endothall thioanhydride inhibits protein phosphatases-1 and -2A in vivo. Am J Physiol 269:C1176–84.

39. Taylor ER, Morgan IM. 2003. A novel technique with enhanced detection and quantitation of HPV-16 E1- and E2-mediated DNA replication. Virology 315:103–109.

40. Das D, Smith NW, Wang X, Richardson SL, Hartman MCT, Morgan IM. 2017. Calcein represses human papillomavirus 16 E1-E2 mediated DNA replication via blocking their binding to the viral origin of replication. Virology 508:180–187.

41. Ohle C, Tesorero R, Schermann G, Dobrev N, Sinning I, Fischer Ts, Adkins NL, Niu H, Sung P, Peterson CL, Aguilera A, GarcÃ-a-Muse T, Arudchandran A, Cerritelli S, Narimatsu S, Itaya M, Shin DY, Shimada Y, Crouch RJ, Belotserkovskii BP, Neil AJ, Saleh SS, Shin JH, Mirkin SM, Hanawalt PC, Blackwood JK, Rzechorzek NJ, Bray SM, Maman JD, Pellegrini L, Robinson NP, Britton S, Dernoncourt E, Delteil C, Froment C, Schiltz O, Salles B, Frit P, Calsou P, Cerritelli SM, Crouch RJ, Cerritelli SM, Frolova EG, Feng C, Grinberg A, Love PE, Crouch RJ, Chan YA, Aristizabal MJ, Lu PY, et al. 2016. Transient RNA-DNA Hybrids Are Required for Efficient Double-Strand Break Repair. Cell 167:1001--1013.e7.

42. Taylor ER, Dornan ES, Boner W, Connolly JA, McNair S, Kannouche P, Lehmann AR, Morgan IM. 2003. The fidelity of HPV16 E1/E2-mediated DNA replication. The Journal of biological chemistry 278:52223–52230.

43. Jenkinson F, Zegerman P. 2022. Roles of phosphatases in eukaryotic DNA replication initiation control. DNA Repair (Amst) 118:103384.

44. Boner W, Taylor ER, Tsirimonaki E, Yamane K, Campo MS, Morgan IM. 2002. A Functional interaction between the human papillomavirus 16 transcription/replication factor E2 and the DNA damage response protein TopBP1. The Journal of biological chemistry 277:22297–22303.

45. Fradet-Turcotte A, Bergeron-Labrecque F, Moody Ca, Lehoux Ml, Laimins La, Archambault J. 2011. Nuclear accumulation of the papillomavirus E1 helicase blocks S-phase progression and triggers an ATM-dependent DNA damage response. Journal of virology 85:8996–9012.

46. Sakakibara N, Mitra R, McBride AA. 2011. The papillomavirus E1 helicase activates a cellular DNA damage response in viral replication foci. Journal of virology 85:8981–8995.

47. Reinson T, Toots M, Kadaja M, Pipitch R, Allik M, Ustav E, Ustav M. 2013. Engagement of the ATR-Dependent DNA Damage Response at the Human Papillomavirus 18 Replication Centers during the Initial Amplification. Journal of virology 87:951–64.

48. Saiada F, Zhang K, Li R. 2021. PIAS1 potentiates the anti-EBV activity of SAMHD1 through SUMOylation. Cell Biosci 11:127.

49. Langsfeld ES, Bodily JM, Laimins LA. 2015. The Deacetylase Sirtuin 1 Regulates Human Papillomavirus Replication by Modulating Histone Acetylation and Recruitment of DNA Damage Factors NBS1 and Rad51 to Viral Genomes. PLoS pathogens 11:e1005181.

50. Nulton TJ, Olex AL, Dozmorov M, Morgan IM, Windle B. 2017. Analysis of the cancer genome atlas sequencing data reveals novel properties of the human papillomavirus 16 genome in head and neck squamous cell carcinoma. Oncotarget 8:17684–17699.

51. Anayannis NV, Schlecht NF, Ben-Dayan M, Smith RV, Belbin TJ, Ow TJ, Blakaj DM, Burk RD, Leonard SM, Woodman CB, Parish JL, Prystowsky MB. 2018. Association of an intact E2 gene with higher HPV viral load, higher viral oncogene expression, and improved clinical outcome in HPV16 positive head and neck squamous cell carcinoma. PloS one 13:e0191581.

52. Wieland A, Patel MR, Cardenas MA, Eberhardt CS, Hudson WH, Obeng RC, Griffith CC, Wang X, Chen ZG, Kissick HT, Saba NF, Ahmed R. 2020. Defining HPV-specific B cell responses in patients with head and neck cancer. Nature doi:10.1038/s41586-020-2931-3.

